# Effects of distraction on taste-related neural processing

**DOI:** 10.1101/693754

**Authors:** I. Duif, J. Wegman, M. Mars, C. de Graaf, P.A.M. Smeets, E. Aarts

## Abstract

Distracted eating is associated with increased food intake and overweight. However, the underlying neurocognitive mechanisms are unknown. To elucidate these mechanisms, 41 healthy normal-weight participants received sips of high- and low-sweet isocaloric chocolate milk, while performing a high- or low-distracting detection task during fMRI on two test days. Subsequently, we measured *ad libitum* food intake. As expected, a region in the primary taste cortex – located in the insula – responded more to the sweeter drink. Distraction did not affect this right insula sweetness response across the group, but did weaken sweetness-related connectivity of this region to a secondary taste region in the right orbitofrontal cortex. Moreover, distraction-related attenuation of taste processing in the insula predicted increased subsequent *ad libitum* food intake after distraction between subjects. These results reveal a previously unknown mechanism explaining how distraction during consumption attenuates neural taste processing and increases food intake. The study was preregistered at https://osf.io/vxdhg/register/5771ca429ad5a1020de2872e?view_only=e3207cd6567f41f0a1505e343a64b5aa.

## Introduction

Worldwide, the prevalence of obesity has nearly tripled since 1975. In 2016, more than 1.9 billion adults were overweight, with 650 million clinically obese (WHO, 2016). The problem of obesity has been partly attributed to the obesogenic food environment, which offers an enormous variety of palatable, energy-dense, easily consumed foods (de Graaf and Kok, 2010; B. J. Rolls, 2010). Furthermore, people’s lifestyles have changed over the last decades, with increasing demands of multi-tasking due to their interaction with electronic devices (e.g. televisions, computers, and smart phones (Carrier, Rosen, Cheever, and Lim, 2015)). As a consequence, people often eat while engaged in activities that prevent them from focusing on satiation signals such as sensory stimulation from the food products they are consuming or gastric signals (e.g. de Graaf and Kok, 2010; Gore, Foster, DiLillo, Kirk, and Smith West, 2003). Such ‘mindless’ or distracted eating has been causally related to increased immediate and later food intake, and is associated with increases in BMI (Bickham, Blood, Walls, Shrier, and Rich, 2013; Bolhuis, Lakemond, de Wijk, Luning, and de Graaf, 2013; de Graaf and Kok, 2010; Grabenhorst and Rolls, 2008; Robinson, Kersbergen, and Higgs, 2014; Wilkie, Standage, Gillison, Cumming, and Katzmarzyk, 2016; Laurson, Lee, and Eisenmann, 2015).

However, the underlying neurocognitive mechanism of how distracted eating could increase food intake, remains elusive. It has been suggested that distraction attenuates taste perception due to limited attentional capacity, which then leads to overconsumption (van der Wal and van Dillen, 2013). However, this putative mechanism has never been tested. An increased understanding of the neurocognitive mechanism could not only reveal the different factors influencing distraction-related overeating, but may also shed light on individual differences in the susceptibility for overeating.

We hypothesize that distraction attenuates processing in the primary and secondary taste cortices, located in the insula and orbitofrontal cortex (OFC) respectively (see e.g. Grabenhorst and Rolls, 2008). The primary taste cortex has been associated with identification, pleasantness and intensity of tastes (Dalenberg, Hoogeveen, Renken, Langers, and ter Horst, 2015; Grabenhorst and Rolls, 2008; Small et al., 2003; Spetter, Smeets, de Graaf, and Viergever, 2010). The OFC receives direct input from the primary taste regions in the insula and has been related to reward-related taste processing, such as hedonic evaluation (E. T. Rolls, Yaxley, and Sienkiewicz, 1990; Small et al., 2007, 2003). Satiety modulates processing in both the primary and secondary taste cortices; both regions show greater taste activation in a state of hunger (Haase, Cerf-Ducastel, and Murphy, 2009; Small, 2010; Small, Zatorre, Dagher, Evans, and Jones-gotman, 2001; Rolls, Sienkiewicx, and Yaxly, 1989). Thus, distraction during food consumption – e.g., due to multi-tasking – might affect processing of primary and higher order taste regions and their connectivity, resulting in attenuated processing of satiety signals and increased food intake. To test this hypothesis, we used a within-subject fMRI design, in which 41 participants performed a high or low distracting categorical visual detection task (**Figure 1)**. Similar to the study by van der Wal and van Dillen (2013), distraction was operationalized by varying the task’s (attentional) load: 90%-high/10%-low load trials on one day (high distraction day) and 90%-low/10%-high load trials on another day (low distraction day). While performing the task in the MR scanner, participants also received sips of isocaloric chocolate milk with a high or low level of sweetness, or a neutral solution. Additionally, we assessed blood glucose and self-reported satiation and satiety on regular time intervals on both the high and low distraction day, as well as *ad libitum* intake of a chocolate snack food at the end of each day.

**Figure 1.**
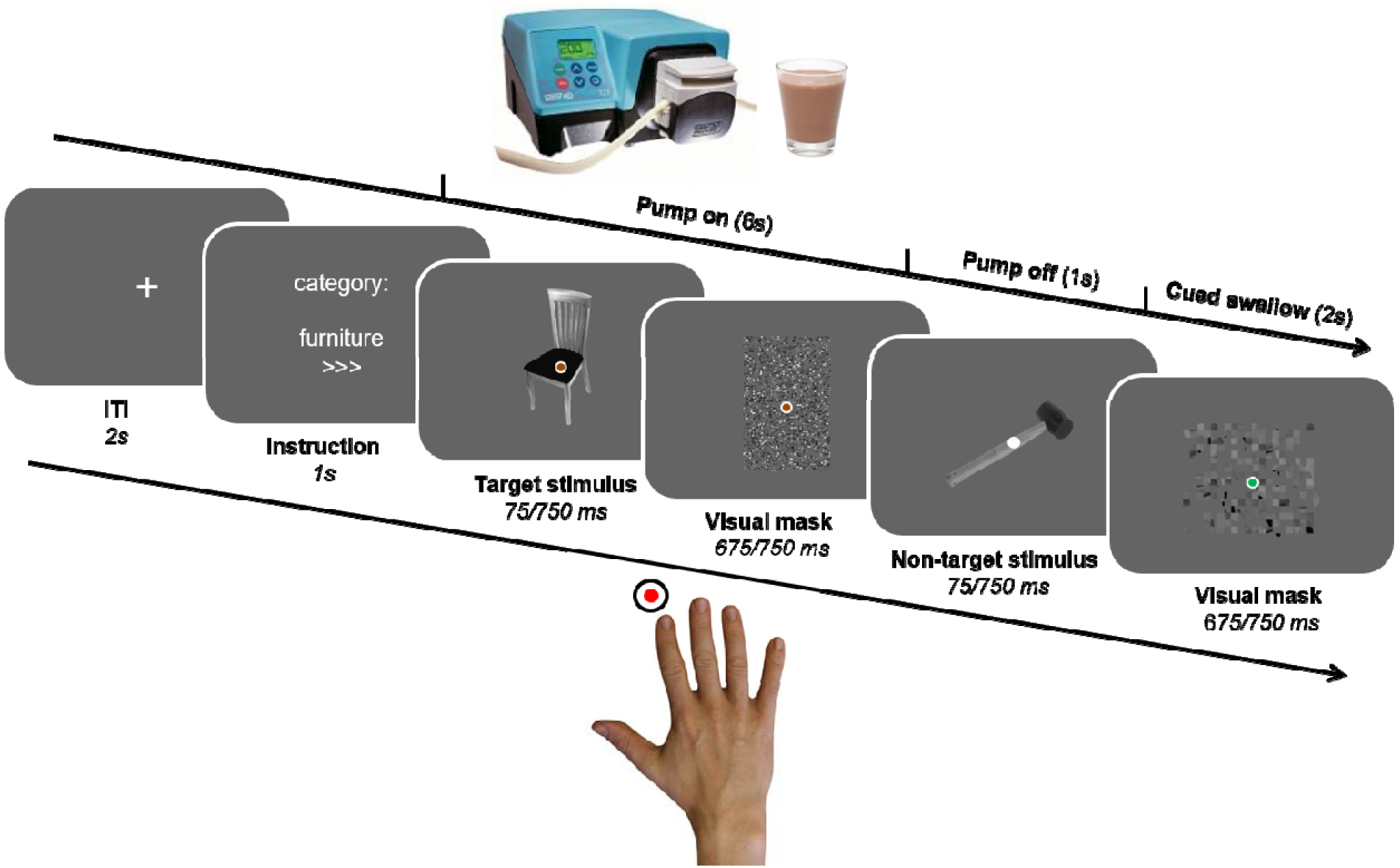
Trial structure of the categorical visual detection task. Each trial started with an instruction screen, indicating the target category (furniture, tool, or toys) and attentional load (low (‘>’) or high (‘>>>’) of the trial. Then, pictures were presented followed by a visual mask, and subjects were instructed to push a button as fast as possible upon detection of pictures belonging to the instructed category. During each trial, subjects received a sip of low or high-sweetened chocolate milk, or a sip of tasteless neutral solution through a gustometer. Markers were placed on the participant’s neck to enable detection of participants’ swallow movements. On- and offsets of the swallow movements were used to determine trial durations in the first level (single-subject) fMRI models.

## Results

### Initial liking of gustatory stimuli

To activate taste-related brain regions, sips of high or low sweet chocolate milk were administered via a gustometer during a categorical visual detection task participants performed in the MR-scanner (see **Figure 1**). Prior to the task, participants rated their liking of the two drinks (**Table S1**) that were selected based on equal liking after a pilot study (see Methods: gustatory stimulation). Results show a marginally significant main effect of Drink Sweetness (high sweet drink, M = 6.1(0.3); low sweet drink, M = 5.5(0.3), *F*(1,34) = 4.53, *p* = .041). At baseline, participants liked the high sweet drink significantly more than the low sweet drink. Therefore, all following fMRI results were corrected for this pre-experimental difference by adding it as a covariate to the analyses. None of the effects below (activation or connectivity) covaried with initial liking (all *p* > .1).

The low sweet drink was perceived equally far from participants’ ideal sweetness as the high sweet drink. For rinsing purposes, participants received a neutral solution in 20% of trials (see *Additional analyses: Liking and ideal sweetness ratings* for the ideal sweetness ratings and liking differences with the other drinks). The neutral trials were not included in the analyses below.

### Performance under distraction

To assess the effects of distraction, we manipulated attentional load of the detection task (**Figure 1**). We tested whether our attentional load manipulation was effective by comparing performance between the two test days. Indeed, participants detected fewer targets when they were rapidly presented, i.e. during the high frequent trials (90% high load trials) on the high distraction day (*d*-prime (± SEM): 2.43 (0.10)) than when they were slowly presented, i.e. during the high frequent trials (90% low load trials) on the low distraction day (*d*-prime (± SEM): 4.00 (0.10), (*F*(1,40) = 264.11, *p* < .001). The sweetness of the chocolate milk administered via the gustometer varied equally across the low- and high-distraction trials: 50% high sweet chocolate milk trials, 50% low sweet chocolate milk trials. As expected, the difference in sweetness did not affect performance (Drink Sweetness (low, high): *F*(1,40) <1, *p* = .748; Drink Sweetness x Load: *F*(1,40) <1, *p* = .907).

### Functional MRI results: effect of distraction on neural taste processing

To determine whether distraction, operationalized as attentional load, affected neural taste processing, we tested the two-way interaction effect of Load (low>high load) and Sweetness (high>low sweetness). Before testing this effect, we first determined whether our load and sweetness manipulations activated the expected brain regions (i.e. fronto-parietal attention network, e.g. Dosenbach et al., 2007, and insula/OFC, respectively).

On our whole-brain corrected threshold (pFWE(cluster-level)<.05), we found effects of attentional load in BOLD responses of a visual and a temporal region when only taking the high frequent trials (90% high load trials versus the 90% low load trials) into account, i.e. when comparing between test days (high>low distraction day). However, when contrasting between the high and low distraction days, while also taking the low frequent trials into account (high + low distraction day: high load trials > low load trials, across drink types), we found areas typically activated in tasks varying in attentional load, including visual and fronto-parietal regions (**Table 1**).

**Table 1.**
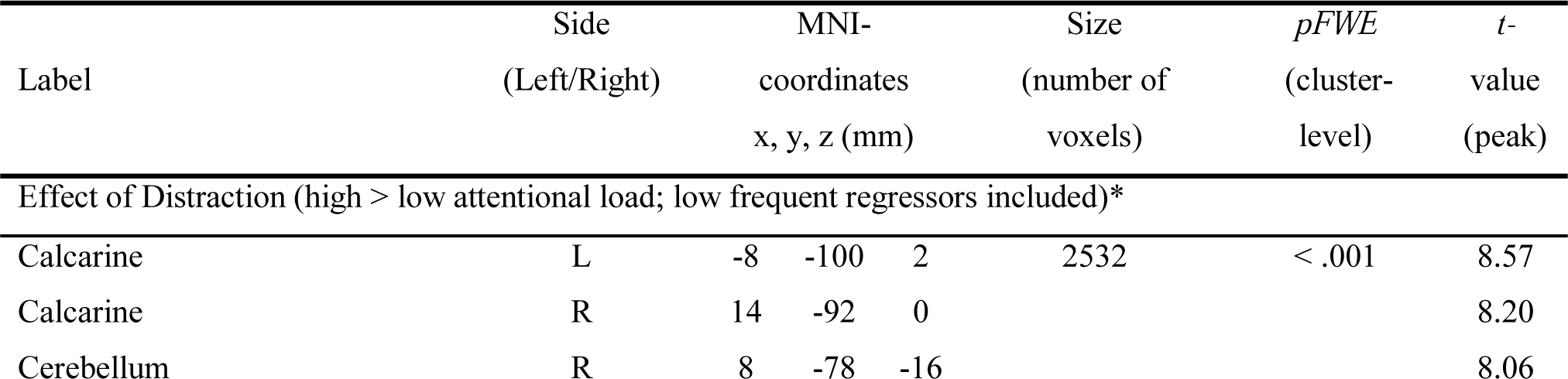
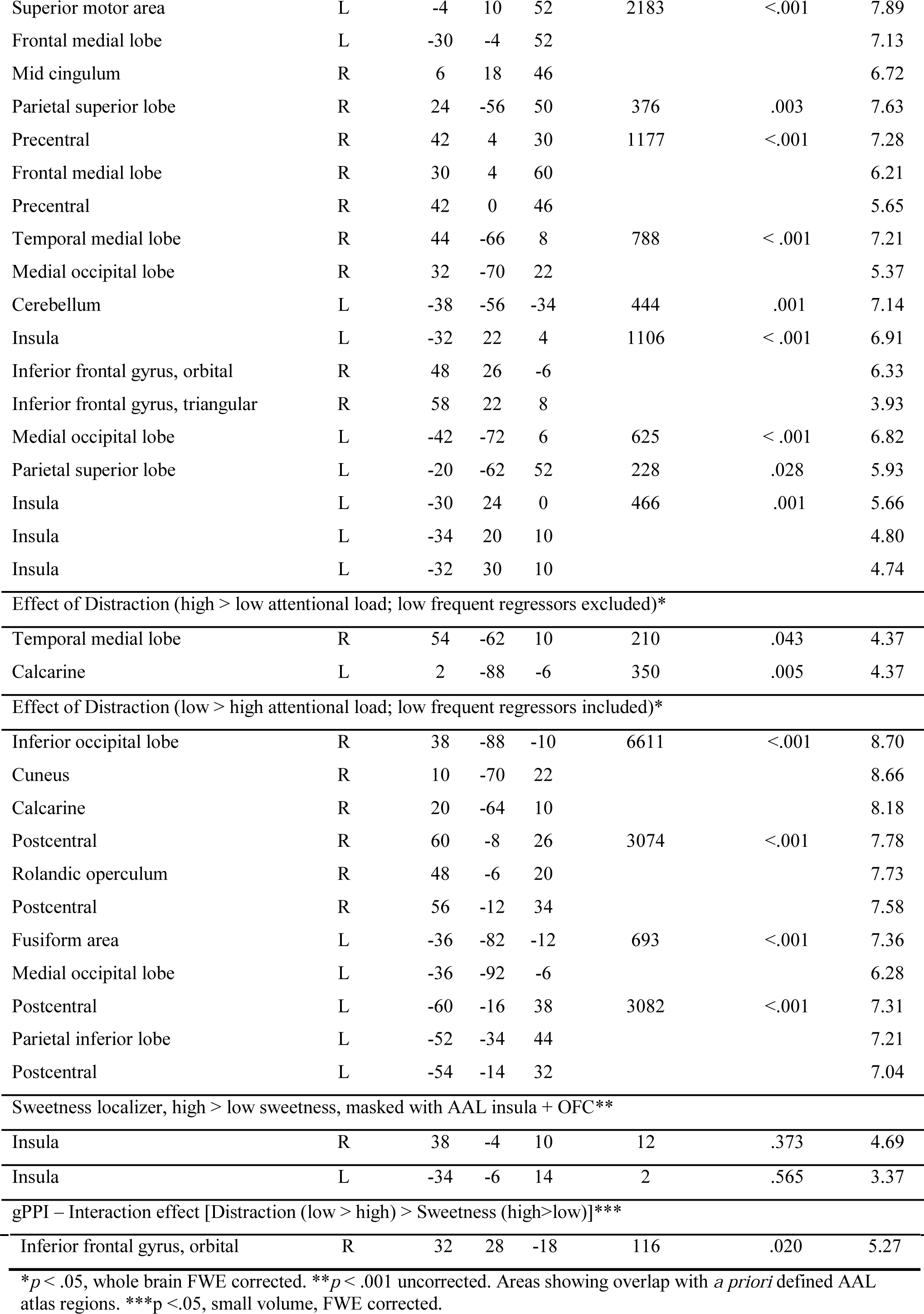
Summary of brain regions exhibiting effects of distraction (attentional load), sweetness, interactions between distraction and sweetness, and the result of the connectivity analysis (gPPI). For the main effects of distraction, results are shown for the comparisons including and excluding the low frequent regressors (see Methods).

| Label | Side<br>(Left/Right) | MNI-coordinates<br>x, y, z (mm) | Size<br>(number of<br>voxels) | $pFWE$<br>(cluster-level) | $t$-<br>value<br>(peak) |
| --- | --- | --- | --- | --- | --- |
| Effect of Distraction (high > low attentional load; low frequent regressors included)* |  |  |  |  |  |
| Calcarine | L | -8 -100 2 | 2532 | < .001 | 8.57 |
| Calcarine | R | 14 -92 0 |  |  | 8.20 |
| Cerebellum | R | 8 -78 -16 |  |  | 8.06 |

|  |  |  |  |  |  |  |  |
| --- | --- | --- | --- | --- | --- | --- | --- |
| Superior motor area | L | -4 | 10 | 52 | 2183 | <.001 | 7.89 |
| Frontal medial lobe | L | -30 | -4 | 52 |  |  | 7.13 |
| Mid cingulum | R | 6 | 18 | 46 |  |  | 6.72 |
| Parietal superior lobe | R | 24 | -56 | 50 | 376 | .003 | 7.63 |
| Precentral | R | 42 | 4 | 30 | 1177 | <.001 | 7.28 |
| Frontal medial lobe | R | 30 | 4 | 60 |  |  | 6.21 |
| Precentral | R | 42 | 0 | 46 |  |  | 5.65 |
| Temporal medial lobe | R | 44 | -66 | 8 | 788 | < .001 | 7.21 |
| Medial occipital lobe | R | 32 | -70 | 22 |  |  | 5.37 |
| Cerebellum | L | -38 | -56 | -34 | 444 | .001 | 7.14 |
| Insula | L | -32 | 22 | 4 | 1106 | < .001 | 6.91 |
| Inferior frontal gyrus, orbital | R | 48 | 26 | -6 |  |  | 6.33 |
| Inferior frontal gyrus, triangular | R | 58 | 22 | 8 |  |  | 3.93 |
| Medial occipital lobe | L | -42 | -72 | 6 | 625 | < .001 | 6.82 |
| Parietal superior lobe | L | -20 | -62 | 52 | 228 | .028 | 5.93 |
| Insula | L | -30 | 24 | 0 | 466 | .001 | 5.66 |
| Insula | L | -34 | 20 | 10 |  |  | 4.80 |
| Insula | L | -32 | 30 | 10 |  |  | 4.74 |
| Effect of Distraction (high > low attentional load; low frequent regressors excluded)* |  |  |  |  |  |  |  |
| Temporal medial lobe | R | 54 | -62 | 10 | 210 | .043 | 4.37 |
| Calcarine | L | 2 | -88 | -6 | 350 | .005 | 4.37 |
| Effect of Distraction (low > high attentional load; low frequent regressors included)* |  |  |  |  |  |  |  |
| Inferior occipital lobe | R | 38 | -88 | -10 | 6611 | <.001 | 8.70 |
| Cuneus | R | 10 | -70 | 22 |  |  | 8.66 |
| Calcarine | R | 20 | -64 | 10 |  |  | 8.18 |
| Postcentral | R | 60 | -8 | 26 | 3074 | <.001 | 7.78 |
| Rolandic operculum | R | 48 | -6 | 20 |  |  | 7.73 |
| Postcentral | R | 56 | -12 | 34 |  |  | 7.58 |
| Fusiform area | L | -36 | -82 | -12 | 693 | <.001 | 7.36 |
| Medial occipital lobe | L | -36 | -92 | -6 |  |  | 6.28 |
| Postcentral | L | -60 | -16 | 38 | 3082 | <.001 | 7.31 |
| Parietal inferior lobe | L | -52 | -34 | 44 |  |  | 7.21 |
| Postcentral | L | -54 | -14 | 32 |  |  | 7.04 |
| Sweetness localizer, high > low sweetness, masked with AAL insula + OFC** |  |  |  |  |  |  |  |
| Insula | R | 38 | -4 | 10 | 12 | .373 | 4.69 |
| Insula | L | -34 | -6 | 14 | 2 | .565 | 3.37 |
| gPPI – Interaction effect [Distraction (low > high) > Sweetness (high>low)]*** |  |  |  |  |  |  |  |
| Inferior frontal gyrus, orbital | R | 32 | 28 | -18 | 116 | .020 | 5.27 |

\* $p < .05$ , whole brain FWE corrected. \*\* $p < .001$ uncorrected. Areas showing overlap with *a priori* defined AAL atlas regions. \*\*\* $p < .05$ , small volume, FWE corrected.

Next, we localized brain regions responding to the difference in sweetness of the chocolate milk (high>low sweetness, p<.001, uncorrected). As expected, within our *a priori* defined search volume, i.e. the anatomically defined bilateral insula + bilateral OFC from the AAL atlas, clusters in the left and right insula responded to this difference (**Table 1** and **Figure 2**). As the left cluster comprised only two voxels, further analyses focused on the right functional insula cluster for the effects of distraction. We did not find differential sweetness responses in the orbitofrontal cortex.

**Figure 2.**
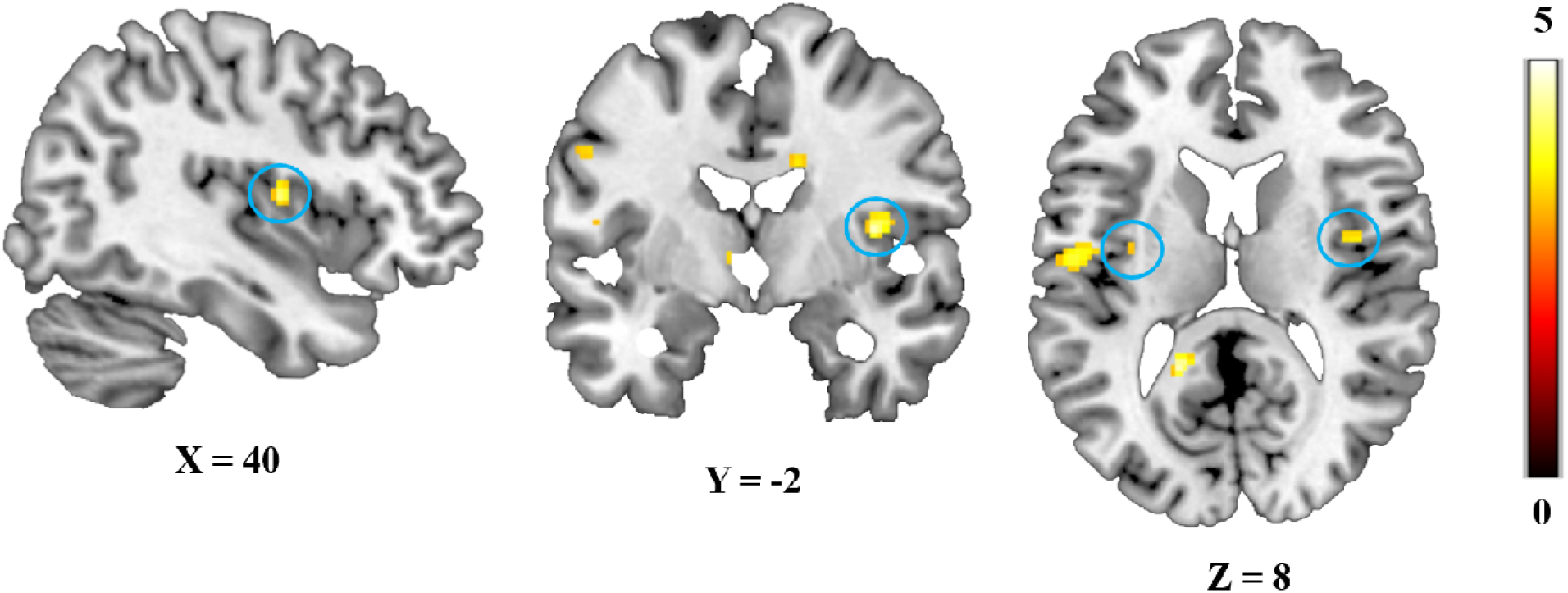
Effect of Sweetness. Response of the left (2 voxels) and right (12 voxels) middle insula to the high vs. low sweet drink. Statistical parametric maps were thresholded at *p* < .001 uncorrected for visualization purposes. All statistical parametric maps were overlaid onto a T1-weighted canonical image. Slice coordinates are defined in MNI152 space and images are shown in neurological convention (left=left).

Finally, we tested the effect of distraction (attentional load) on activation of the right insula cluster that responded to the sweetness manipulation. We found no evidence for this effect, as the averaged extracted condition parameter estimates across the right insula cluster did not show a Load x Sweetness interaction (*F*(1,40) <1, *p* = .711). We obtained similar results when session order (whether participants had the low distraction day first or second) was added to the analysis as between-subject factor. However, when we used this right insula cluster that showed greater responses for high>low sweetness (at *p*<.001, uncorrected) as a seed in a secondary functional connectivity (i.e. generalized psychophysiological interaction) analysis, it showed decreased connectivity with a region in the right OFC under high compared with low distraction during processing of sweet taste (high>low sweetness, **Table 1** and **Figure 3**). This right OFC region ([32,28,-18], AAL: right inferior frontal gyrus, pars orbitalis]) was significant within our *a priori* defined search volume of the insula plus OFC, (*pFWE*(cluster, after small volume correction (SVC)) = .020, *t* = 5.27, k = 116). This shows that distraction weakens functional connectivity between the right insula and right OFC during taste processing.

**Figure 3.**
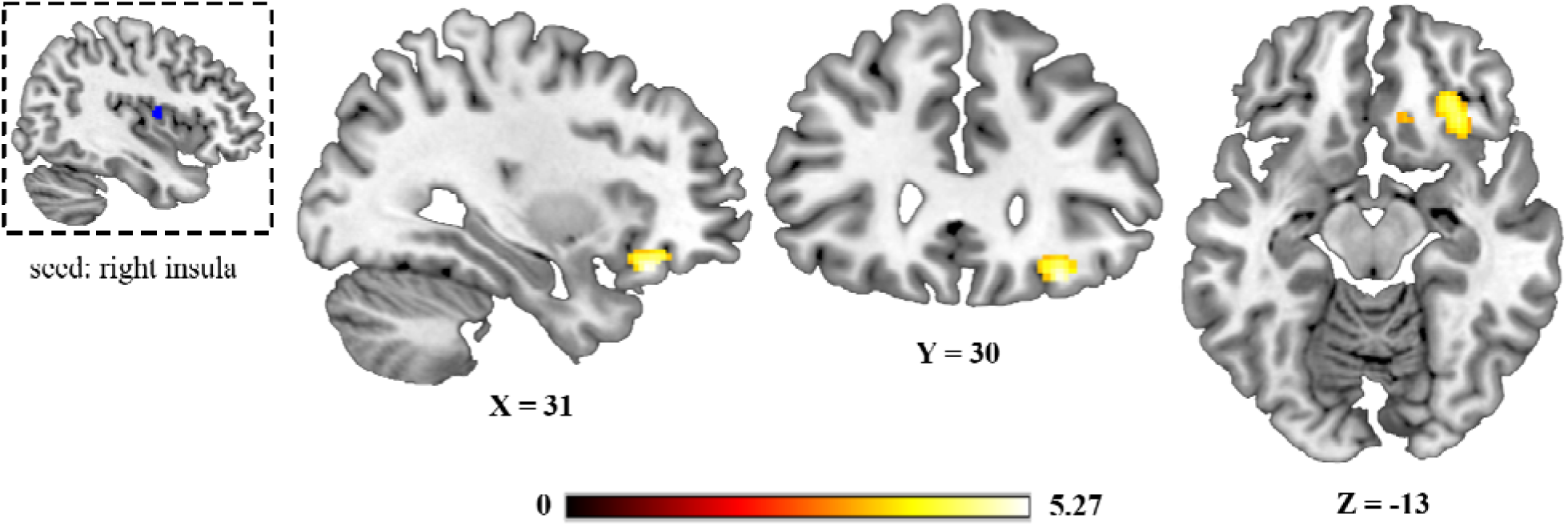
Results of the gPPI analysis with the right insula seed region in the top left (in blue, extracted from the high>low sweetness contrast). Shown is the right orbitofrontal area exhibiting significantly (*p* = .020 after SVC) higher sweetness-related functional connectivity with the seed region under low, relative to high, distraction.

### Effect of distraction on self-reported satiety: hunger, fullness, and thirst ratings

Hunger, fullness, and thirst ratings that were filled out digitally during the task (t_0_, t_5_, t_10_, t_30_) and on paper before and after the test day (t_-5_, t_75_) were analyzed separately (**Table 2**). All ratings showed main effects of time, except for hunger, which was not significant in the paper ratings. These effects indicate significant increases in fullness and significant decreases in thirst and hunger over the time course of the task (digital) or test day (paper), as anticipated. No effects of distraction, i.e. attentional load, on any of the ratings were found. In addition, we found no significant pre-experimental differences between the low and high distraction day for all self-reported measures. Although distraction did not affect self-reported satiety ratings, we did observe distraction-related differences when correlating it with consumption-induced increases in blood glucose, as indicated below.

**Table 2.** Self-report satiety, filled out digitally and on paper, averaged over distraction (high, low attentional load). Means and standard errors per time point, and Time statistics.

| | $t_{(-5)}$ | $t_{(0)}$ | $t_{(5)}$ | $t_{(10)}$ | $t_{(30)}$ | $t_{(75)}$ | p | F |
| --- | --- | --- | --- | --- | --- | --- | --- | --- |
| *Digital satiety ratings* |  |  |  |  |  |  |  |  |
| Hunger | - | 6.8(0.2) | 6.4(0.2) | 5.9(0.3) | 5.9(0.3) | - | .005 | 5.068 |
| Fullness | - | 2.2(0.2) | 2.9(0.3) | 3.7(0.4) | 4.4(0.4) | - | <.001 | 11.210 |
| Thirst | - | 6.0(0.3) | 4.2(0.4) | 3.6(0.4) | 3.8(0.4) | - | .001 | 15.189 |
| *Satiety ratings on paper* |  |  |  |  |  |  |  |  |
| Hunger | 6.3(0.2) | - | - | - | - | 6.4(0.2) | ns. | 2.049 |
| Fullness | 2.3(0.2) | - | - | - | - | 3.6(0.3) | <.001 | 20.617 |
| Thirst | 5.3(0.3) | - | - | - | - | 4.3(0.3) | .004 | 9.503 |

### Effect of distraction on blood glucose levels

We collected blood at four time points during the experiment (finger prick method; t_0,_ t_30,_ t_50,_ and t_75_) to assess the effects of distraction on glucose level increase during and after the computer-paced chocolate milk intake. Analysis of the blood glucose levels revealed a main effect of Time, *t*(1, 88.85) = 11.28, *p* < .001, **Figure 4**), meaning that participants’ blood glucose levels increased significantly over the four time points during and after chocolate milk consumption, as expected (M_t0_ = 4.43 (0.46), M_t30_ = 4.84 (0.74), M_t50_ = 7.14 (1.23), M_t75_ = 7.17 (1.29)). Furthermore, distraction (attentional load) tended to affect these glucose increases over time, with reduced increases in glucose levels on the high, relative to the low, distraction day (*t*(1, 260.19) = 1.81, *p* = .072). This effect was driven by the significant distraction-related decreased rise in glucose levels at t = 75 relative to baseline (low distraction day: M_t75-t0_ = 2.94(1.48), high distraction day: M_t75-t0_ = 2.50(1.32), *t*(1, 113.96) = 2.09, *p* = .039). Interestingly, this distraction-induced attenuation of blood glucose rise correlated negatively with changes in hunger ratings (*t*(1, 37) = −3.44, *r* = -.50, *p* = .002). This correlation was driven by a significant effect on the low distraction day, on which increases in blood glucose rise were related to decreased hunger ratings (*t*(1, 37) = −2.14, *r* = −.34, *p* = .039). On the high distraction day, there was no correlation between hunger and glucose (*t*(1, 37) = 0.56, *r* = .09, *p =* .581). Thus, distraction tended to decrease the glucose response to the consumed chocolate milk and attenuated the association between glucose and self-reported hunger.

**Figure 4.**
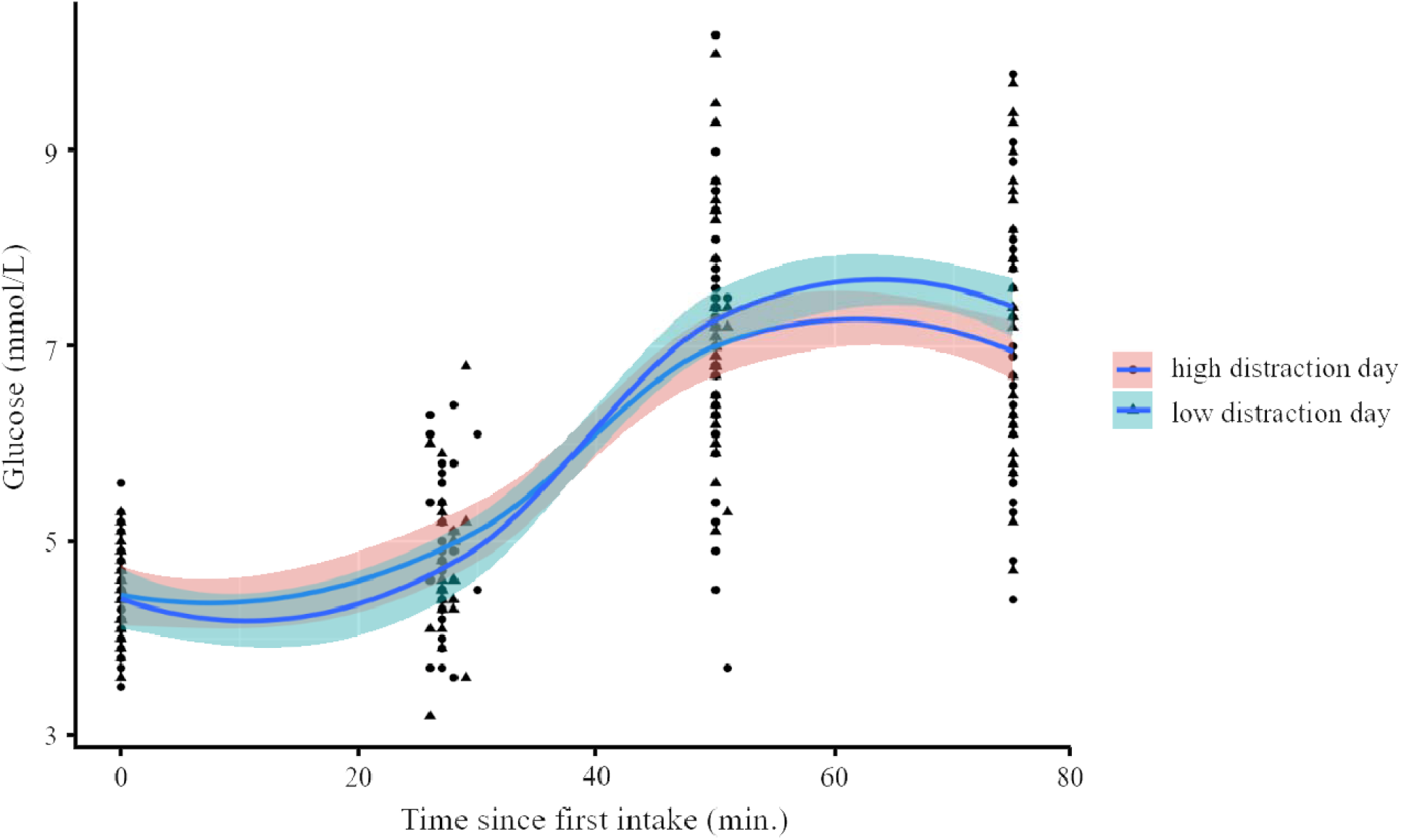
Blood glucose response to the chocolate milk per session (high, low distraction day) for each time point (at baseline (0 g. of chocolate milk consumed), right after the task (240 g. consumed), 50 and 75 minutes after baseline) in mmol/L. Blood glucose increases were marginally lower on the high versus low distraction day. Loess lines of best fit were used to fit the data.

### Effect of distraction on food intake

Forty-five minutes after completing the distraction task in the MR-scanner, we determined whether distraction (attentional load) during earlier chocolate milk consumption affected the total amount of chocolate snacks consumed *ad libitum*. Chocolate snack intake after the scan session did not differ between the test days (M_lowload_ = 65.6(5.9), M_highload_ = 68.1(6.8), *t*(1,40) = −0.61, *p* = .546). However, further analyses showed a significant interaction between attentional load and session order (low distraction day first, high distraction day first) for snack intake (*F*(1,39) = 8.27, *p* = .007). Food intake was significantly higher on the second, relative to the first, test day, independent of the attentional load of the test day (M_testday1_ = 61.2(6.2) g., M_testday2_ = 72.4(6.4) g., *F*(1,40) = 8.68, *p* = .005, **Figure S1A**). This result is in line with results of a pilot experiment in which 31 participants drank chocolate milk *ad libitum* 45 minutes after computer-paced chocolate milk consumption while performing a more or less demanding visual detection task (see Additional analyses: *Behavioral pilot study*). In the pilot study, we found similar results, i.e. participants consumed more chocolate milk on the second test day than on the first (M_testday1_ = 73.2(10.3) g., M_testday2_ = 111.8(19.0) g., *F*(1,21) = 8.24, *p* = .008, **Figure S1B**). It is therefore likely that the interaction between load and session order for food intake is driven by a repetition effect. Given session order effects on food intake, session order was added as a between-subject factor to all other analyses. None of other reported results changed after correction for order, or interacted with order (all *p* > .1).

### Distraction-related brain-behavior correlations

Above, we showed that – across the group – our distraction manipulation diminished insula-OFC connectivity, but did not affect insula sweetness responses, later food intake, or self-reported satiety measures. The above-mentioned glucose-hunger association as a function of distraction already demonstrated that there is variability between participants in how they respond to consumption under distraction. To further explore inter-individual differences in the fMRI data, we assessed whether the interaction effect (Load x Sweetness) in the right insula area co-varied with the effects of distraction on the behavioral outcome measures (self-reported hunger and fullness, food intake) and blood glucose levels. Interestingly, right insula activation during the fMRI task co-varied with *ad libitum* intake of the chocolate snack 45 minutes after completing the task (Load x Sweetness x food intake: *F*(1,39) = 4.81, *p* = .023, *r* = .36). Subsequent analyses showed this relation was present on the high (*F*(1,39) = 9.61, *r* = .45, *p* = .004), but not on the low distraction day (*F*(1,39) <1, *r* = −.01, *p* = .973). More specifically, only activation for the low sweet drink on the high distraction day tended to predict how much participants would subsequently eat on the high distraction day after fMRI (low sweet drink: *F*(1,39) = 3.87, *r* = −.30, *p* = .056, high sweet drink: *F*(1,39) <1, *r* = −.04, *p* = .823). Thus, individuals in which high attentional load attenuated insula activation of the low sweet drink, showed increased subsequent food intake (**Figure 5**).

**Figure 5.**
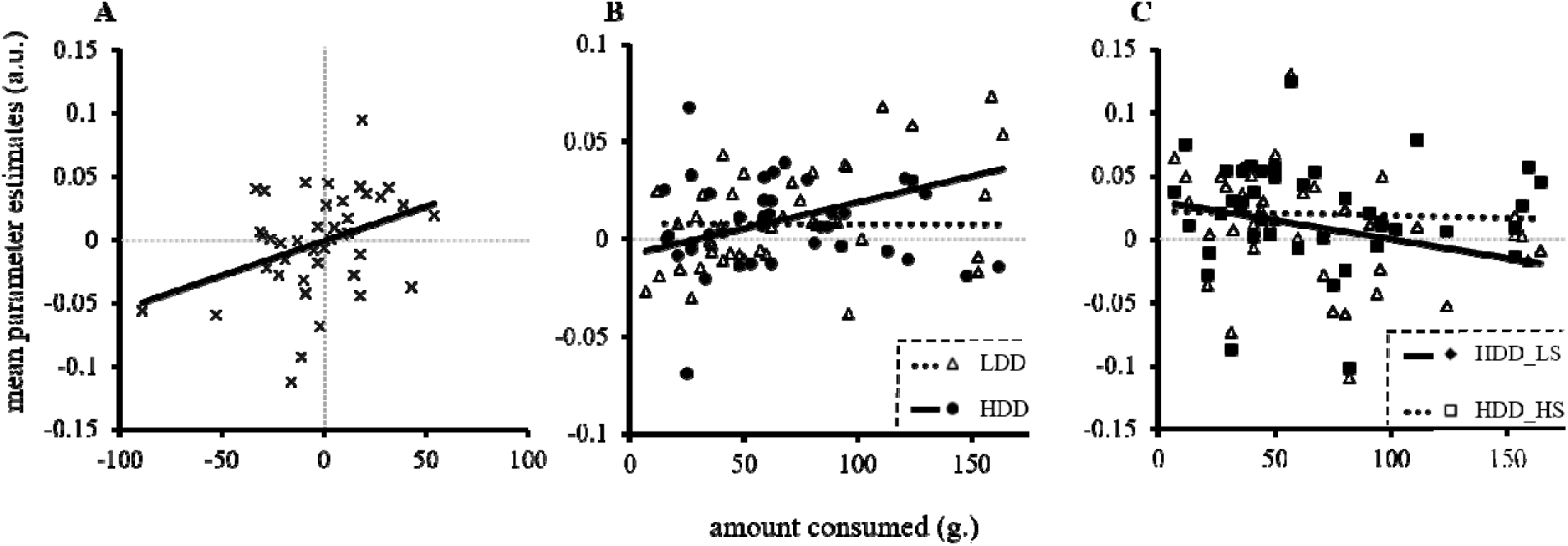
Brain-behavior correlations for the relation between taste-related (high>low sweetness) responses in the right insula, distraction (high, low attentional Load) and *ad libitum* food intake. Panel A) significant brain-behavior correlation at the highest level (Load (low-high distraction day) x Sweetness (high-low sweet drink) x *ad libitum* intake (low-high distraction day), *r* = .36, *p* = .023. Panel B) separate correlations for the low (LDD, *r* = -.01, *p* = .973) and high (HDD) distraction day (*r* = .45, *p* = .004, high-low sweetness. Panel C) correlations on the high distraction day only, for low (LS, *r* = -.30, *p* = .056) and high (HS, *r* = -.04, *p* = .823) sweetness separately. Less activation for the low, but not the high, sweet drink on the high (but not low) distraction day seems to predict increased food intake on the high distraction day. Mean parameter estimates are presented in arbitrary units (a.u.), *ad libitum* food intake in amount consumed in grams.

Including BMI and performance on the task as covariates did not change the above-reported pattern of findings. Finally, we did not find correlations for brain activation in the right insula and blood glucose levels, hunger, or fullness ratings, or any brain-behavior correlations for insula-OFC connectivity.

### Additional results

#### Liking and ideal sweetness ratings

We expected liking ratings to decrease significantly for the low and high sweet, but not the neutral, drink over the time course of the task in the MR scanner as a result of sensory-specific satiety. Results show a main effect of Time (*F*(1,19) = 6.49, *p* = .003)), indeed reflecting significant decreases in liking after the task compared to baseline for the low and high sweet drinks, but not for the neutral drink (see **Table S1** for means and standard errors / deviations, and statistics). Explorative analysis of the effect of distraction (attentional load) on liking ratings showed no significant results (interaction effect of Load, Drink Type and Time: *F*(1,16) <1, *p* = .795). At baseline, participants liked the high sweet drink significantly more than neutral drink (high sweet drink, M = 6.1(0.3); neutral drink, M = 4.5(0.5), *t*(1,35) = 2.64, *p* = .012). There was a marginally significant difference in liking ratings between the low sweet and neutral drink (low sweet drink, M = 5.4(0.3); neutral drink, M = 4.5(0.5), *t*(1,34) = 1.73, *p* = .092).

Furthermore, we assessed whether participants rated both the low and high sweet drink equally far from the optimum (a rating of 5) in terms of how well the drinks matched their ideal sweetness. As expected, we found no significant differences at baseline on this measure for the low relative to the high sweet drink, showing that the low sweet drink was perceived equally far from participants’ ideal sweetness as the high sweet drink (mean difference from optimum: high sweet drink, M = 1.5(0.2); low sweet drink, M = 1.0(0.2), *t*(1,26) = 1.47, *p* = .153). There were no significant decreases on these ratings over time *F*(1,14) = 1.42, *p* = .280, **Table S1**), nor as a function of distraction *F*(1,14) <1, *p* = .430).

#### Imagined desire for something sweet or savory

A repeated measures ANOVA with within-subject factors Load (low, high), Taste (sweet, savory) and Time (t_0_, t_30_) revealed significant main effects of Taste, Time, and an interaction effect of Taste and Time on participants’ imagined desire for something sweet or savory (**Table S1**). The main effect of Taste reflects a larger overall desire for “something savory” relative to “something sweet” (*F*(1,37) = 92.45, *p* < .001), and the main effect of Time indicates a significant overall reduction in desire (*F*(1,37) = 23.51, *p* < .001). Finally, the significant Taste x Time interaction reflected a larger decrease in desire for the sweet taste over time relative to the savory taste, showing successful induction of sensory specific satiety for the sweet taste (*F*(1,37) = 17.16, *p* < .001).

## Discussion

Distracted eating has been convincingly associated with increased food intake (Robinson et al., 2013), but the underlying neurocognitive mechanisms remained elusive. Here, we demonstrated how distraction – operationalized by varying attentional load – affects taste activation in, and connectivity between, primary and secondary taste-related brain areas (insula and OFC respectively), blood glucose levels, subsequent chocolate snack intake and self-reported satiation and satiety.

As expected, high compared with low sweetness elicited differential BOLD responses in the insula: a small (2-voxel) cluster in the left insula and a larger cluster (12 voxels) in the right middle insula. These findings fit with recent work by Dalenberg et al. (2015), who showed that the insular cortex processes the presence, pleasantness, and concentration of taste. More specifically, the right insular cortex dominated processing of taste concentration (intensity) signals, whereas the left insular cortex was more involved in representation of the presence of a taste stimulus and its pleasantness. This is in line with results from two other studies that also showed lateralization of intensity to the right insula (Small et al., 2003; Spetter et al., 2010). Some studies have related activity of the right insula to processing of pleasantness, however, they did not control for effects of intensity (Nitschke et al., 2006; Small et al., 2001). In the present study, we varied taste intensity by contrasting activation in response to a high sweet versus a low sweet drink, and corrected for subjective liking differences of the two drinks at baseline. Crucially, this correction did not change the results of the high>low sweetness contrast. Therefore, the currently observed responses of the insula in the high>low sweetness comparison were mainly driven by differences in intensity, explaining the dominance of the right insula.

Importantly, we demonstrated that distraction attenuated taste-related functional connectivity between the right insula – found for the high>low sweetness contrast – and an area in the OFC. This OFC region is located in the caudomedial OFC (cmOFC, Small et al., 2007), which is thought to be a relay between the anterior insula and the caudolateral OFC (clOFC) which is responsive to the pleasantness of taste (Small et al., 2007). Another study manipulated pleasantness of chocolate milk and tomato juice through satiation, and showed that pleasantness of the drinks correlated with taste activation in the left and right OFC, with the latter overlapping with the region found in our study (Kringelbach, O’Doherty, Rolls, and Andrews, 2001). Thus, our findings suggest diminished functional coupling between primary and secondary taste cortices by distraction.

Our results further indicate that some individuals were more sensitive to distraction-related attenuation of taste-related activation in the right insula than others. Only when high attentional load affected taste processing of the low sweet drink in the insula, subsequent food intake of participants increased. Previous work also showed large variation between participants in the effects of distraction on food intake (Bellisle, Dalix, Airinei, Hercberg, and Péneau, 2009; Martin, Coulon, Markward, Greenway, and Anton, 2009). Furthermore, the meta-analysis by Robinson et al. (2013) showed that specifically highly disinhibited eaters are less likely to decrease their food intake after or during distraction. The inter-individual variability found in our study is therefore not surprising and future studies should further investigate what drives these individual differences.

One study also investigated effects of (working memory) load on food-related processing during fMRI (van Dillen and van Steenbergen, 2018). In that study, higher cognitive load diminished nucleus accumbens responses during categorization of high-versus low-calorie food pictures. In addition, they showed that cognitive load altered the functional coupling between the nucleus accumbens and the right dorsolateral prefrontal cortex for high-versus low-calorie food pictures (van Dillen and van Steenbergen, 2018). However, these authors studied cognitive load effects on hedonic brain responses in the nucleus accumbens during categorization of high- and low-calorie food pictures versus object pictures as edible or inedible, in the absence of consumption during the task. By assessing actual taste processing during consumption of drinks in the scanner, we add to these previous findings that distraction attenuates connectivity in the taste network, and that the attenuating effect of attentional load on taste-related processing in the insula predicts subsequent food intake.

We found that responses for the low sweet drink in particular predicted increases in later food intake under high load. Interestingly, a study by Hoffmann-Hensel, Sijben, Rodriguez-Raecke, and Freiherr (2017) found a similar effect when assessing the impact of cognitive load on fMRI responses to low and high calorie food odors with the same working memory task of varying load by van der Wal and van Dillen (2013). Their behavioral data revealed diminished perceived intensity for low-, but not high-, calorie food odors during high cognitive load. Similarly, higher cognitive load decreased OFC responses for low-calorie, but not the high-calorie, food odors. Their results could be explained by higher saliency of the high-calorie food odors and this may apply to our findings as well; as the high sweet drink is more salient compared with the low sweet drink due to its higher sweetness, attentional load might less easily suppress high sweet taste perception. It is, however, important to note that olfaction and gustation follow similar, but not identical neural pathways (e.g. Hoffmann-Hensel et al., 2017; Small et al., 1999). Nevertheless, the current and previous findings suggest that the saliency of odors and tastes plays a role in how distraction affects processing of food-related stimuli.

We observed a marginally significant effect of distraction on blood glucose levels, i.e. the increase in glucose levels was reduced after high (versus low) distraction. This was driven by a significant distraction-related decreased rise in glucose levels when comparing blood glucose levels 45 minutes after chocolate milk consumption (t_75_) to baseline (t_0_). The effect of distraction on glucose levels correlated negatively with its effect on self-reported hunger. These findings should be interpreted with caution, as the effect of attentional load on glucose did not relate to subsequent food intake. Therefore, we cannot infer that lower blood glucose levels are related to increased consumption, despite the negative correlation with hunger. Rather than *ad libitum* consumption at a fixed time point, glucose declines have been associated with earlier meal initiation (Melanson, Westerterp-Plantenga, Saris, Smith, and Campfield, 2017; Strubbe and Woods, 2004). Our results point towards lower glucose levels at t_75_ after high (relative to low) distraction, which could have resulted in participants initiating a subsequent meal sooner. However, we did not test voluntary meal initiation in this study and future studies should assess how distraction-related differences in glucose levels affect this.

One limitation of our study is that the design was optimized for the primary outcome measure, i.e. the fMRI effects. Therefore, the distraction manipulation had to be relatively subtle. Distractions such as watching television versus doing nothing in the MR-scanner provide a less-controlled fMRI comparison with more noise than varying attentional load. The relative subtleness of the distraction manipulation could explain why we did not find group effects of attentional load on food intake, or on self-reported satiety measures.

In conclusion, by using fMRI during consumption, we found that distraction reduced functional connectivity between taste processing areas and that distraction-related attenuation of taste-related processing in the insula predicted subsequent food intake. This provides a neurocognitive mechanism that improves our understanding of (the susceptibility for) overeating, and points to an important role for undisrupted taste processing in overeating. A better understanding is essential for successful prevention and treatment of overweight and obesity, where being mindful about the taste of food during consumption could be part of the solution. In our current – overly distracting – society, attentive eating might be more important than ever, to protect taste processing from being disrupted. Future studies should investigate the role of attention in neural taste processing in obesity.

## Acknowledgements

This work was supported by a public-private Food, Cognition, and Behavior grant funded by the Netherlands Organization for Scientific Research (NWO, grant 057-14-001) with financial support of Mars Wrigley. We would like to thank Eva Ketel, Marlou Lasschuijt, Lieke van Lieshout and Natalia Staniszewska for their help in conducting the study.

## Author Contributions

Conceptualization: E.A., J.W., P.S., M.M., C.G., and I.D.; Methodology: E.A., J.W., P.S., M.M., C.G., and I.D.; Software: J.W. and I.D.; Formal Analysis: J.W., I.D, and E.A.; Investigation: J.W. and I.D.; Writing – Original Draft: I.D., E.A.; Writing – Review & Editing: E.A., J.W., P.S., M.M., and C.G.; Visualization: J.W., I.D. and E.A.; Supervision: E.A. and J.W.; Project Administration: E.A., P.S., and J.W.; Funding Acquisition: J.W., E.A., C.G., and P.S.

## Methods

### Participants

Forty-six right-handed healthy adults, who were recruited from Nijmegen and surroundings through advertisement, participated in the study. They gave written informed consent and were reimbursed for participation according to institutional guidelines of the local ethics committee (CMO region Arnhem-Nijmegen, the Netherlands, 2015-1928).

As a result of drop-out (i.e. not completing the second test day (n = 1), technical problems (n = 4)), the final sample size of the study was 41 (age range 18 – 35 y; mean age 22.5 y (3.5); 31 females; mean (SD) Body Mass Index (BMI) 21.9 kg/m^2^ (1.89); mean (SD) waist-to-hip ratio (WHR): 0.80 (0.05)).

### Screening

During an intake, the study was explained to the participant, commitment and availability of the participant was checked, and physical measurements (BMI (weight(kg)/(height (cm^2^)) and waist-hip ratio (waist(cm)/hip(cm)) were measured. The participant practiced the task (see below) to avoid between-session effects, and filled out questionnaires to screen for inclusion and exclusion criteria. To be eligible for participation in the study, participants had to have a BMI within a range of 18.5 – 30.0, had to be within 18 – 35 years old, and right-handed. Exclusion criteria were current pregnancy; MRI-incompatibility; diabetes mellitus; history of hepatic, cardiac, respiratory, renal, cerebrovascular, endocrine, metabolic or pulmonary diseases; uncontrolled hypertension; neurological, psychiatric, or eating disorders; current strict dieting; restrained eating score ≥ 3.60 for females and ≥ 4.00 for males on the Dutch Eating Behavior Questionnaire (DEBQ, van Strien et al., 1986); current psychological or dietary treatment; taste or smell impairments; use of neuroleptica or other psychotropic medication; food allergies relevant to the study, deafness, blindness, and sensori-motor handicaps; drug, alcohol or nicotine addiction; inadequate command of both Dutch and English, and a change in body weight of more than 5 kg in the past two months.

### Procedure

Participants were invited to the laboratory for two experimental test days: a low and a high distraction day. The order was randomized: half of the participants had the low distraction day first and the high distraction day second, the other half of participants had the inverse order. Prior to each test day, participants were instructed to abstain from eating solid foods and from drinking sugared or sweetened drinks (but not water) three hours prior to the experiment, and to refrain from alcohol use (24 hours) and drug use (7 days). Participants were provided with a standardized meal (yogurt drink, strawberry flavor (Breaker, Melkunie, Nijkerk, the Netherlands), which they were instructed to eat three hours before each test day.

At the start of the first test day, anthropometric measurements were taken (weight and waist-hip ratio). Furthermore, participants rated how hungry, full, and thirsty they felt on a paper version of a 100mm visual analogue scale (VAS) ranging from 0 (“not at all”) to 10 (“very”). Subsequently, participants underwent an fMRI scan for 30 minutes in which they performed a categorical visual detection task (**Figure 1**). Before, during (after the first and second task block), and directly after the task, participants rated their hunger, fullness, and thirst again, now on digital VASs. Before and after the task, participants also digitally rated how nauseous and anxious they felt, and their desire for something savory and something sweet. During the task, participants received chocolate milk through small tubes. After scanning, participants watched a documentary (BBC Life, Primates or Plants, order was randomized). After the documentary, participants rated how hungry, full, and thirsty they were once more on paper. Subsequently, participants were seated in front of a bowl with *ad libitum* colored button-shaped chocolates (M&Ms, Mars Wrigley) for ten minutes, and were asked to eat until comfortably full. At test day 1, the participants completed the Behavioural Inhibition System / Behaviour Approach System (BIS/BAS; Carver and White, 1994), Baratt Impulsiveness Scale-11 (BIS-11; Patton, Stanford, and Barratt, 1995), and Kirby (delayed reward discounting; Kirby, 2009) questionnaires. At test day 2, they completed the following questionnaires: Binge Eating Scale (BES; Gormally, Black, Daston, and Rardin, 1982), Food Frequency Questionnaire – Dutch Healthy Diet (FFQ-DHD; van Lee et al., 2016), DEBQ (Strien and Frijters, JER, 1986), and Power of Food Scale (PFS; Lowe et al., 2009), which are not taken into consideration for the current analyses.

During the test day, glucose measurements were taken through finger pricks and analyzed with use of a glucose meter (Stat Strip Xpress®, Nova Biomedical, Waltham, MA). This was done at four time points, t_0_ (baseline, before first chocolate milk exposure); t_30_ (directly following last exposure to chocolate milk), t_50,_ and t_75_ (before consumption of the chocolate snack), to assess whether the distraction manipulation affected blood glucose levels.

At least one week after their first test day (mean difference (± SD): 11.93 (7.92) days), participants revisited the lab for their second test day, at the same time of day when possible (mean time difference (± SD: 1.00 (± 1.52) hrs.

### Gustatory stimulation

To avoid differences in liking between the low and high sweet chocolate drinks participants received during the task in the MR-scanner, we performed a pilot study in which 7 solutions of cocoa powder (Blooker, 2 g.), dextrine-maltose (Fantomalt, 9 g.) in whole milk (3.5% fat/100 g.), and liquid artificial non-caloric sweetener (Natrena, ranging from 0.0867 to 1.955 g., in steps of 0.2669) were rated by 10 participants (who did not participate in the current study) on a 100mm visual analogue scale (VAS), labeled “not at all liked” and “very much liked”. Using the average liking rating from this pilot study, the concentrations for low and high sweet drinks were determined such that both the low and high intensity drink were at equal distance from the optimum. The two concentrations chosen for the low and high sweet chocolate milk contained 0.0867 and 1.5457 g. of Natrena per 100 g. whole milk.

A similar approach was used to determine the neutral solution. Based on previous work by van Veldhuizen et al. (2010), we created 4 solutions containing 2.5 mM sodium bicarbonate and 25 mM potassium chloride in water. We also created three weaker versions at 25%, 50% and 75% of the original concentration. The most neutral concentration (closest to a liking rating of 5 on a scale of 0 to 10) was used, which was the solution at 50% of the original concentration.

Before and after the task participants performed in the MR-scanner, they received 2 rounds of sips (3.75 mL) of the neutral drink and of the low and high sweet drink for tasting through tubes innervated by pumps (gustometer, Watson-Marlow, Ontario). For all drinks, they rated how much they liked the drink on VASs. For the two chocolate milk drinks, they also rated to what extent the sweetness of the chocolate milk matched their ideal sweetness on an ideal point scale.

### Categorical Visual Detection Task

Participants performed a categorical visual detection task during 3T fMRI scanning (**Figure 1**). Each trial (total duration: 12s) started with a fixation cross (duration 2s), followed by an instruction screen (1s), indicating the category of pictures to which the participant needed to respond (furniture, tools, or toys), and the speed of the trial (‘>’ for a slow trial, ‘>>>’ for a fast trial. For example, if the instruction screen stated: category: furniture, >>>”, this meant participants needed to respond to stimuli in the category furniture, and the pictures would be presented at high speed. In order to keep visual stimulation equal for both trial types, a visual mask always followed a picture. The visual masks were scrambled versions of the stimulus pictures, to keep luminance equal. For the low speed trials, both pictures and visual masks were presented for 750ms. For the high speed trials, pictures were presented for 75ms, and the visual mask for 675ms. Consequently, there were twice as many pictures and visual masks in the high speed trials relative to the low speed trials (12 vs. 6), thus, a higher attentional load. Whenever a target stimulus was presented (i.e. a picture belonging to the instructed category), participants had to push a button upon detection with their right index finger. Participants made responses using an MRI-compatible button box. Participants received no feedback on whether they responded correctly.

During the trials, participants received a sip (3.75 mL) of chocolate milk of high or low sweetness, or a sip of the neutral solution through a gustometer. Drink administration started along with presentation of the first picture, and lasted for 6 seconds. A dot changing in color in the center of the screen informed participants of the start (brown color) and finish (white color, 1s) of administration. At the end of each trial, the dot turned green (2s), cueing participants to swallow the sip. As swallowing can also be an uncontrollable, reflexive movement, participants’ swallowing was filmed. A marker was placed on the Adam’s apple, as this area shows the most swallow-related movement (**Figure 1**). Frame-by-frame video analysis of the marker’s movement was later performed to pinpoint the exact moments in time when participants swallowed during the experiment.

Participants performed four blocks of 20 trials (a total of 80 trials). For the low distraction day, 90% of the trials were low-speed trials (pictures presented at a slow pace), and 10% of the trials were high-speed trials (pictures presented at a fast pace). Thus, on the low distraction day, each block contained 18 low-speed trials, and 2 high-speed trials. For the high distraction day, this division was the same, however in the opposite direction (90% high difficulty trials, 10% low difficulty trials). Each block had four neutral trials. Trials 1, 7, 14 and 20 were always neutral. Of the remaining trials, 50% were of high sweetness, the other 50% of low sweetness. Category and drink sweetness presentation were pseudo-randomized, i.e. the same category and sweetness were never presented more than 3 times in a row. Moreover, maximally two target stimuli were presented after another.

### Behavioral analyses: initial liking of gustatory stimuli

To test for pre-experimental differences in liking of the high compared to low sweet chocolate milk we performed a repeated-measures ANOVA with within-subject factor Drink Sweetness (low sweet, high sweet) on the mean baseline ratings.

### Behavioral analyses: performance

We used the sensitivity index d-prime (*d’*) to calculate participants’ task performance. From the task, four types of response were obtained: hits (target was detected correctly), miss (target was present, but the participant incorrectly indicated there was no target), false alarms (participant indicated a target was present when there was not), and correct rejections (participant correctly indicated there was no target). D-prime was calculated using the formula: *d’* = Z_Hit_ – Z_FA_ (Haatveit et al., 2010; Snodgrass and Macmillan, 1990), where “Hit” represents the proportion of hits when a target was present (hits/(hits + misses)), also known as the hit rate, and “FA” represents the proportion of false alarms when a target was absent (false alarms/(false alarms + correct rejections)), the false-alarm rate. D-prime is then calculated by taking the difference between the Z-transforms of these two rates. The Z-transformation was done using the statistical formula NORMSINV(Hit)– NORMSINV(FA) in Matlab (2016a). To avoid *d’* scores reaching -∞ or +∞, perfect scores were adjusted by subtracting 0.0025 from the hit rate, and adding 0.0025 to the false alarm rate. This correction resulted in maximum *d’*-scores of +5.61 (100% hits, 0% false alarms), and minimum scores of −5.61 (0% hits, 100% FA).

Mean d-prime scores on the detection task were analyzed using repeated measures ANOVA (IBM SPSS Statistics 23, Chicago, IL) with attentional Load (low, high) and Drink Type (low, high, neutral) as within-subject factors. Low frequent conditions (i.e. 10% low speed trials on the high distraction day and vice versa) were excluded from this analysis, as the low number of trials in these conditions would likely bias the results.

### Imaging and fMRI analyses

#### (f)MRI Data Acquisition

To measure blood oxygen level dependent (BOLD) contrast, whole-brain functional images were acquired on a Siemens 3T Skyra MRI scanner (Siemens Medical system, Erlangen, Germany) using a 32-channel coil. During the task, 3D echo planar imaging (EPI) scans using a T2*weighted gradient echo multi-echo sequence (Poser, Versluis, Hoogduin, and Norris, 2006) were acquired (voxel size 3.5 × 3.5 × 3 mm isotropic, TR = 2070 ms, TE = 9 ms; 19.25 ms; 29.5 ms; 39.75 ms, FoV = 224mm). The slab positioning and rotation (average angle of 14 degrees to AC axis) optimally covered both prefrontal and deep brain regions. Before the acquisition of functional images, a high-resolution anatomical scan was acquired (T1-weighted MPRAGE, voxel size 1×1×1 mm, TR 2300 ms, TE 3.03 ms, 192 sagittal slices, flip angle 8°, field of view 256 mm).

#### (f)MRI Image Processing

Data were analyzed using SPM8 (www.fil.ion.ucl.ac.uk/spm) and FSL version 5.0.11 (http://www.fmrib.ox.ac.uk/fsl/). The volumes for each echo time were realigned to correct for motion artefacts (estimation of the realignment parameters is done for the first echo and then copied to the other echoes). The four echo images were combined into a single MR volume based on 30 volumes acquired before the actual experiment started using an optimized echo weighting method (Poser et al., 2006). Combined functional images were slice-time corrected by realigning the time-series for each voxel temporally to acquisition of the middle slice and spatially smoothed using an isotropic 8 mm full-width at half-maximum Gaussian kernel. Next, ICA-AROMA (Pruim et al., 2015) was used to reduce motion-induced signal variations in the fMRI data. Subject-specific structural and functional data were then coregistered to a standard structural or functional stereotactic space (Montreal Neurological Institute (MNI) template) respectively. After segmentation of the structural images using a unified segmentation approach, structural images were spatially coregistered to the mean of the functional images. The resulting transformation matrix of the segmentation was then used to normalize the anatomical and functional images into Montreal Neurological Institute space. The functional images were resampled at voxel size 2 × 2 × 2.

#### Video analysis of swallow movements

Recordings of participants’ swallow movements were made with an in-bore camera and infrared LED light (MRC Systems GMbH). Matlab software (Matlab, version 2016a) was used to detect the circle-shaped marker placed on the participant’s neck in the video. After detection, a rectangular frame-of-interest was determined around the marker using the subject-specific center coordinates and radius of the marker to reduce the search area. To determine the size of the frame of interest, the x- and y-coordinates of the marker’s center and its radius were multiplied by two. Next, the marker’s radius was added to or subtracted from the center coordinates to calculate the boundaries of the frame of interest in the –x, +x, −y and +y direction. To time lock the video data to the MR scanner pulses, a beep was recorded at the onset of the first scanner pulse. Subsequently, frame-to-frame video intensity differences were extracted from the videos and coupled with the onsets of trials to determine when participants swallowed the chocolate milk. To detect on- an offsets of the swallows, the Hilbert transform was used, a function that can determine the envelope of a waveform in an analytical signal (<u>Matlab</u> version 2016a).

#### Statistical fMRI analysis

Statistical analysis of fMRI data was performed using a general linear model (GLM) approach. The images of both experimental runs were combined into one model including the low and high distraction test days. At the individual (first) level, subject-specific data were analyzed using a fixed effects model, which included five regressors of interest. The first four reflected the trials of low attentional load, low sweet drink; low attentional load, high sweet drink; high attentional load, low sweet drink; and high attentional load, high sweet drink. The fifth regressor reflected trials in which the neutral solution was given, which was always of high frequent load, so of low attentional load on the low distraction day, and of high attentional load on the high distraction day to rinse in between chocolate milk trials. Durations reflected the moment the gustometer started drink administration until the first swallow of the participants, or in case this video data was not available, the moment the swallow was cued in the task (dot turning green, mean duration: 11.22s (SD: 0.80s/ SEM: 0.01s). All regressors were convolved with the canonical hemodynamic response function. Parametric modulators reflecting the number of button presses per trial were added to the model for each regressor of interest, to correct for signal change induced by the difference in number of targets between the low and high attentional load condition. High pass filtering (128s) was applied to the time series of the functional images to remove low-frequency drifts and correction for serial correlations was done using an autoregressive AR(1) model. Signal variation in white matter and cerebrospinal fluid regions was also included.

At the group (second) level, we assessed the effect of Load (high > low attentional load) in two ways. First, we contrasted high with low load trials by including the high frequent trials of each test day only, meaning a contrast between sessions (high distraction day: high load regressors across drink types > low distraction day: low load regressors across drink types). Second, we also added the low frequent regressors to assess this contrast (high + low distraction day: high load regressors across drink types > low load regressors across drink types).

Taste-related brain areas (sensitive to sweetness) were localized with the contrast high > low Sweetness (*p*<.001, uncorrected), over both test days. For this contrast, within-test day data was available; therefore, we did not make a second contrast including low frequent regressors. We used the Automated Anatomical Labeling (AAL) atlas (Tzourio-Mazoyer et al., 2002) to determine whether the areas activated in this functional contrast overlapped with the anatomical insula (bilateral insula) and OFC (bilateral superior, medial, mid, and inferior orbitofrontal regions of this atlas.

To investigate the effect of attentional Load on processing in primary and secondary taste-related areas, we assessed the interaction effect of Load (high > low Load, using the test days in which these conditions were the high-frequent Load condition) x Sweetness (high>low). The activated taste-related regions (determined by the contrast high > low sweetness) were used as regions-of-interest (ROIs) for the Load x Sweetness interaction contrast. Mean beta weights were extracted from all voxels in both ROIs separately using MarsBar (Brett, Anton, Valabregue, and Poline, 2002). The regionally averaged beta-weights were analyzed using ANOVA with the same factors as in the whole-brain analyses. As two ROIs were tested, effects were considered significant when reaching a threshold of p<.025 (Bonferroni-corrected for multiple comparisons). The results of all random effects fMRI analyses were thresholded at *p* < 0.001 (uncorrected) and statistical inference was performed at the cluster level, family-wise-error-corrected (P_FWE_<0.05) for multiple comparisons over the search volume (the whole brain). All statistical parametric maps were overlaid onto a T1-weighted canonical image. Slice coordinates are defined in MNI152 space and images are shown in neurological convention (left=left).

#### Distraction-related functional connectivity analysis

As a secondary analysis, we performed a generalized psychophysiological interaction (gPPI; McLaren, Ries, Xu, and Johnson, 2012) analysis to investigate distraction-related differences in functional connectivity for taste-related processing. As a seed, we used a taste-related region from our localizer approach described above. To estimate the neural activity producing the physiological effect in the seed region for each subject, the BOLD signal was extracted from this region and deconvolved (Gitelman, Penny, Ashburner, and Friston, 2003). This was included in the model as the physiological regressor, as were the durations for each of the relevant task conditions (low load, low sweet drink; low load, high sweet drink; high load, low sweet drink; and high load, high sweet drink), and the psychophysiological interaction was entered by multiplying the estimated neural activity by the duration times for each of the task conditions separately convolved with the HRF, resulting in nine regressors of interest on the first level (i.e., one physiological, four psychological, and four interaction regressors). For each subject, we created a PPI contrast for the interaction effect of distraction (high>low attentional load) and sweetness (high>low sweetness). On the second level, this PPI contrast was analyzed separately using a one-sample t-test. Statistical inference (pFWE<0.05) was performed at the cluster-level, correcting for multiple comparisons over the a priori defined small search volume: bilateral insula and OFC (AAL atlas; (Tzourio-Mazoyer et al., 2002)). The intensity threshold necessary to determine the cluster-level threshold was set at p<0.001, uncorrected.

#### Analyses of secondary outcome measures: food intake, blood glucose levels and self-report satiety ratings

The effect of attentional Load on subsequent *ad libitum* food intake was tested with a paired-samples t-test, to compare the total amount of chocolate snacks consumed (g.) between the high and low distraction test days.

Blood glucose levels were analyzed using the linear mixed model (lmer4, version 1.1-14, Bates, Mächler, Bolker, and Walker, 2014) package in R (version 3.5.1, https://www.r-project.org), because we also expected variation over participants in the glucose response over time, and traditional repeated measures ANOVA cannot account for such variation. Participants’ glucose levels were analyzed with Time (t = 0, t = 30, t = 50 and t = 75) and Load (low, high) as fixed factors. We used random intercepts for participants, as we expected fasted glucose levels to vary between participants. Moreover, we used random slopes for the Time predictor, to account for the expected variation over participants in glucose response over time.

To test whether mean self-report satiety ratings (hunger, fullness, and thirst) varied as a function of Load, we used repeated measures ANOVA with within-subject factors Load and Time (digital: t_0_, t_5_, t_10_, t_30_, paper: t_-5_, t_75_).

#### Analyses: brain-behavior correlations

Exploratory, we investigated whether *ad libitum* food intake, blood glucose levels, and self-report satiety measures (hunger, fullness, thirst; filled out digitally or on paper) covaried significantly with the effect of Load on processing in the taste-related areas differentially activated in the high>low sweetness contrast. For *ad libitum* food intake, a repeated measures ANCOVA was executed with Load (high, low) and Sweetness (high, low) as within-subject factors, and the difference in food intake between the low and high distraction day as covariate. For the blood glucose measurements, the difference in increase over time (t_75_-t_0_) between the low and high distraction day was used a covariate, as this difference drove the effect of load on blood glucose levels described previously. For the hunger, fullness, and thirst digital and paper ratings, the difference over time (digital: t_30_-t_0_, paper: t_75_-t-_5_) between the low and high distraction day was used for the covariates.

#### Additional analyses

##### Liking and ideal sweetness ratings

To test for pre-experimental differences in liking of the three drinks we performed a repeated-measures ANOVA with within-subject factor Drink Type (low sweet, high sweet, neutral) on the mean baseline ratings. Furthermore, we performed a repeated-measures ANOVA with within-subject factors Load (low, high), Drink Type (low sweet, high sweet, neutral), and Time (t_0(1)_, t_0(2)_, t_30(1)_, t_30(2)_) to test whether liking decreased significantly over time for the drinks. Moreover, we exploratively assessed whether liking ratings were affected by attentional load.

With respect to the ratings on how well the low and high sweet drinks matched participants’ ideal sweetness, we aimed to show that both the low and high sweet drinks were at equal distance from the optimum. To test this, we calculated the absolute difference from the optimum by subtracting the optimum (a rating of 5) from the low and high sweet ratings. Subsequently, a paired samples t-test was used to test whether these mean ratings were significantly different. Exploratory, we assessed whether there were changes in these ratings over time, or as a function of attentional load.

##### Imagined desire for something sweet or savory

Before and after the task in the MR scanner, we asked participants how much they desired “something sweet” and “something savory”. If sensory specific satiety was successfully induced, we expected their imagined desire for something sweet, but not savory, to decrease significantly during the task. To test this, we executed a repeated measures ANOVA with within-subject factors Load (low, high), Taste (sweet, savory) and Time (t_0_, t_30_) and assessed the interaction effect between Taste and Time to test whether participants’ satiety decreased specifically for the sweet taste.

##### Behavioral pilot study

Prior to the current study, we performed a behavioral pilot study with a similar set-up as the current study in 31 participants. They performed the same visual detection task on two separate test days (high, low distraction) in an MRI-like set-up in a behavioral lab. However, no actual fMRI scanning was performed. To mimic the MRI-set-up, participants lay on a table and heard MRI-sounds through headphones during the experiment. Participants performed 80 trials on the visual detection task during computer-paced consumption of chocolate milk; however, there were no trials during which the neutral solution was administered via the gustometer. Participants performed 40 trials in the high and low sweetness condition, instead of 32 in the current study. As a result, participants consumed 150 grams of each chocolate milk during the task, instead of 120 grams. Eighty (instead of 90) percent of trials were of high frequent load, and 20% (instead of 10%) of low frequent load. Instead of a chocolate snack, subjects consumed chocolate milk *ad libitum*. No blood glucose measurements were taken.

---

**Table S1.** Self-reported liking, ideal sweetness, and desire for something sweet or savory ratings, averaged over Load (high, low). Related to “Results: Initial liking of gustatory stimuli”. Means and standard errors per time point, and Time statistics.

|  | t<sub>(0)</sub> | t<sub>(5)</sub> | t<sub>(10)</sub> | t<sub>(30)</sub> | p | F |
| --- | --- | --- | --- | --- | --- | --- |
| *Liking ratings* |  |  |  |  |  |  |
| Low sweet drink | 5.4(0.3) | 4.6(0.3) | 3.6(0.3) | 3.9(0.3) | <.001 | 13.92 |
| High sweet drink | 6.1(0.3) | 5.9(0.3) | 5.2(0.4) | 5.1(0.4) | .025 | 3.58 |
| Neutral solution | 4.5(0.5) | 4.6(0.5) | 4.7(0.5) | 4.8(0.5) | .639 | <1 |
| *Ideal sweetness ratings* |  |  |  |  |  |  |
| Low sweet drink | 3.9(0.2) | 3.8(0.3) | 3.5(0.3) | 3.7(0.3) | .364 | 1.37 |
| High sweet drink | 6.4(0.3) | 6.6(0.3) | 6.6(0.3) | 6.6(0.3) | .878 | <1 |
| *Desire for something sweet or savory* |  |  |  |  |  |  |
| Sweet | 5.6(0.3) | - | - | 3.7(0.3) | <.001 | 36.26 |
| Savory | 7.2(0.2) | - | - | 7.0(0.3) | ns. | <1 |

**Figure S1.**
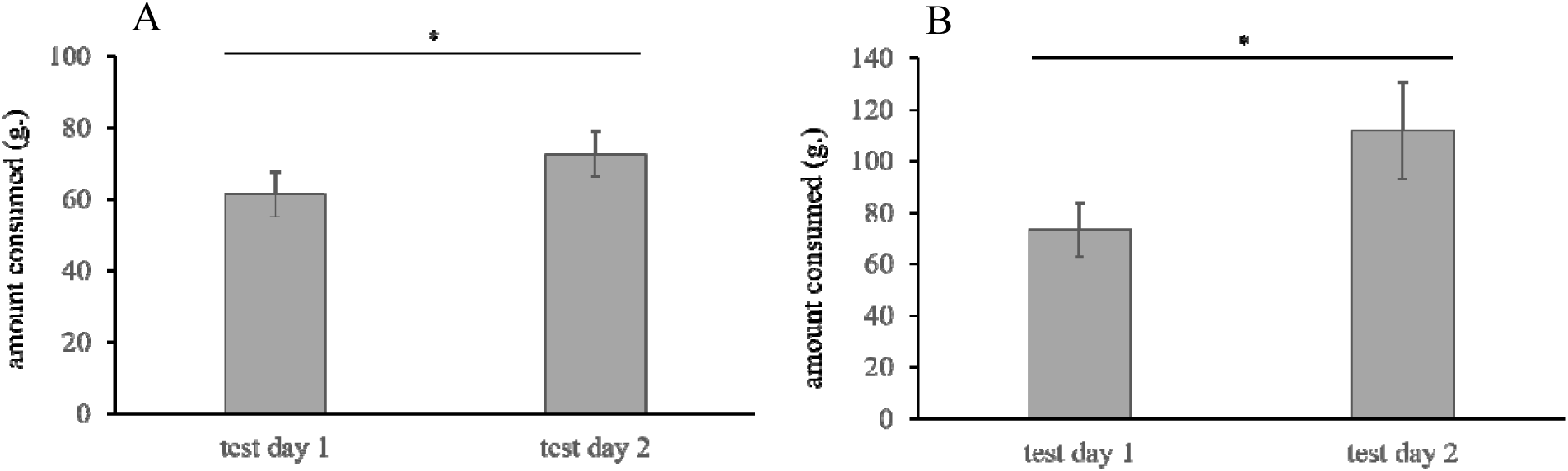
*Ad libitum* intake of the chocolate snack (current study) or milk (pilot study) for each test day (test day 1, test day 2), independent of Distraction (attentional load). Related to “Results: Effect of distraction on food intake”. Error bars depict standard error of the mean. Panel A) Intake in the current study. The mean amount consumed (g.) was significantly higher on the second test day. The asterisk indicates *p* = .008. Panel B) Intake of the chocolate milk in the pilot study. In line with the results of the current study, the mean amount consumed (g.) was significantly higher on the second test day. The asterisk indicates *p* = .005.

